# Evaluation of search-enabled Pre-trained Large Language Models on retrieval tasks for the PubChem Database

**DOI:** 10.1101/2024.08.15.608120

**Authors:** Ash Sze, Soha Hassoun

## Abstract

Databases are indispensable in biological and biomedical research, hosting vast amounts of structured and unstructured data, facilitating the organization, retrieval, and analysis of complex data. Database access, however, remains a manual, tedious, and sometimes overwhelming, task. We investigate in this study the current state of using pre-trained, search-enabled LLMs for data retrieval from biological databases. Equipped with internet search and code generation capabilities, LLMs promise to streamline database access through natural language, expedite search and knowledge retrieval, and provide coherent analytical summaries. As an example database, we focus on evaluating a current search-enabled LLMs (GPT-4o) for retrieval from the PubChem database, a flagship, heavily used database that plays a critical role in biological and biomedical research. As PubChem is an open archival repository, it provides a well-documented programmatic interface that can be exploited through LLM code generation capabilities. We evaluate retrieval tasks for eight common PubChem access protocols that were previously documented. The tasks include identifying interacting genes and proteins, finding drug-like compounds based on structural similarity, retrieving bioactivity data, and locating stereoisomers and isotopomers. We develop a methodology for adopting the protocols into an LLM-prompt, where we supplement the prompt with additional context through iterative prompt refinement as needed. To further evaluate the LLM capabilities, we instruct the LLM to perform the retrieval with and without using programmatic access. We compare the results (referred to as gold and silver answers) when using these retrieval modalities with two traditional retrieval baselines that include running the manual search steps for each reference protocol through the PubChem database web interface, and through the provided PUG (Power-User Gateway) programmatic access. We quantitatively and qualitatively summarize our results, showing that generating programmatic access is more likely to yield the correct answers. We highlight the value and limitations of using current search-based LLMs for database retrieval. We also provide guidance for the future development that can improve the accuracy and reliability of search-based LLMs.

## 1. Introduction

Efficient and instantaneous access to comprehensive and well-organized biological databases is crucial for advancing research, development, and innovation [1]. There are now a multitude of such databases, including PubChem, UniProt, PDB, MetaCyc, BRENDA, KEGG, and many others. Despite the tremendous growth in the size and scope of enzymatic and biological databases, finding and linking relevant data within and across such databases is a daunting task that hinders biological discovery and the development of data-driven machine-learning approaches. While the development of programmatic access facilitates this access, it often requires a programming background, which may hinder access to unskilled biological or biomedical researchers. Importantly, the availability of Large Language Models (LLMs) has the potential to play a transformative role in accessing databases. LLMs, which are built using transformer models[2], tokenize their inputs and learn model parameters based on a training corpus. These models can be used for generative AI: to produce content (text, code, images, and recently video) based on input prompts in natural language.

Such generative capabilities have enabled assistive technologies such as GitHub CoPilot for code generation, Grammarly as a virtual writing assistant, Vi Trainer as a virtual health coach, and Microsoft CoPilot to assist with productivity tasks. For databases, LLMs are poised to simplify and streamline access through a natural language interface and provide coherent analytical summaries, thus expediting search and knowledge retrieval. Indeed, while there are tremendous efforts for using LLMs for Information Retrieval (IR) through re-writing queries and expedited retrieval and ranking of results [3], there are currently limited studies on using pre-trained LLMs such as GPT-4 for database access or on creating customized LLM-based agents for database access. For example, a recent benchmark, BIRD (a BIgbench for laRge-scale Database) with text-to-SQL tasks spanning 95 databases and 37 professional domains, was evaluated using GPT-4 [4]. A recent specialized GPT-3-based LLM is trained on several database related tasks including query rewriting and index tuning [5]. GeneRAG enhances LLMs such as GPT-4 with gene related tasks by retrieval augmentation generation [6]. There is preliminary data supporting that the use of long-context LLMs can even subsume retrieval, RAG, and SQL[7].

We investigate in this paper the use of pretrained search-enabled LLMs for data retrieval from the PubChem database. Search-enabled LLMs are recent models that provide the capabilities of internet searching to provide the LLM with the most up-to-date relevant context. For example, OpenAI’s GPT-4 and GPT-4o (“o” is for omni), a faster version of GPT-4, and Google’s Gemini, provide such capabilities. Hence, such models can provide a relevant specific context for a user query. Importantly, a user could instruct such an LLM to primarily rely on data retrieved from a particular database. Here, we select GPT-4o to explore its capabilities in retrieving data from the PubChem database.

Using GPT-4o, we explore retrieval from the PubChem database. PubChem is a flagship public database at the NCBI of the National Library of Medicine (NLM) [8–13]. PubChem aggregates chemical information from over 995 sources including chemical vendors, research and governmental organization, journal publishers, NIH initiatives such as Pharos [14], and journal publications and information is organized into “data collections”. For example, the Substances are chemical entities uploaded by PubChem contributors while Compounds encompass unique chemical structures extracted, aggregated, and standardized from contributed Substance records. BioAssay records are uploaded from contributing organizations and individuals and contain bioactivity and toxicity data collected through experimentations. The Pathway collection integrates data from multiple pathway databases including the Reactome knowledgebase [15], BioCyc [16], Wikipathways [17] and others. The Protein and Gene collections contain records that have associated biological activities in the BioAssays, or that are found in the Pathway collection. PubChem is therefore effectively an “open” archival system, with freely accessible, searchable, downloadable data and where users can contribute, under clear guidelines, their chemical substances and biological experimental results. Due to its rich content and its scale, PubChem plays a central role in biomedical and life science research, with millions of monthly users [12]. To access its content, PubChem offers programmatic access to its data through Power User Gateway (PUG) REST or CGI services. For example, PUG-View retrieves individual PubChem records, while PUG-REST retrieves specific information from within one or more PubChem Records. PubChem also offers programmatic access to its RDF (Resource Description Framework), where each RDF statement consists of a subject, predicate, and an object. For example, with “soltamox may treat breast cancer”, the subject is soltamox, the object is breast cancer, and the predicate is “may treat”. SPARQL queries that consist of a set of triple patterns in which the subject, predicate and/or object) can be a wildcard can then be used to query the RDF for possible matches.

While effective, programmatic access to PubChem and other databases requires detailed guidance. To facilitate access to PubChem, a recent paper [18] provided a detailed guide on how to effectively use PubChem’s extensive resources and tools for various research purposes. By offering detailed step-by-step instructions, the paper enhances users’ ability to search for and analyze chemical information, perform similarity searches, retrieve bioactivity data, and understand the relationships between chemicals and biological entities. For example, the following six steps are outlined in protocol #8 for retrieving stereoisomers and isotopomers of a compound (valsartan, CID 60846) through identity search:

1. *Navigate to the PubChem homepage; type CID 60846 structure*.
2. *Once the search results appear, click on the “Identity” tab to access identity search results. Then, click on the “Settings” button to adjust the identity search parameters*.
3. *Configure Search Options: The default setting is “Same Stereo Isotope,” which searches for compounds with the same connectivity, stereochemistry, and isotopism. To find stereoisomers, change the setting to “Same Isotope.”*
4. *Download Stereoisomers in CSV format*
5. *Select the “Same Stereo” option to find isotopomers, and*
6. *Download the returned isotopomers in a CSV format*.

This example demonstrates that users may find navigating the extensive interface overwhelming, especially with numerous tabs and sections that need to be explored to locate specific information. The vast amount of data retrieved during searches requires further filtering and interpretation to find relevant results. Additionally, setting up effective search parameters, particularly for advanced searches involving chemical structures and similarity scores, demands intimate knowledge of the search parameters. Finally, regularly updating and managing large datasets derived from PubChem for Machine Learning applications can be cumbersome. These challenges are not unique to PubChem, search and programmatic access for other databases such as the KEGG or UniProt databases can also be challenging.

Here, we explore the use of GPT-4o on commonly used PubChem tasks [18]. The eight tasks include finding genes and proteins that interact with a query compound, finding compounds with various similarity metrics or properties based on 2-D or 3-D similarity searches, computing similarity searches between compounds, retrieving bioactivity data from a substructure search, finding drugs that target a particular gene, finding compounds with classifications, and retrieving a query compound’s stereoisomers and isotopomers. For example, instead of manually executing the steps in protocol #8, the prompt can be, “Based only on information from PubChem, find stereoisomers and isotopomers of valsartan.” Ideally, the results retrieved via the protocol steps and the prompt should be identical. Our evaluation first generates gold and silver answers for each task. The gold answer is obtained via executing the protocol through PubChem’s interface, while the silver answer is generated from executing the programmatic access for that protocol. In some cases, there are differences between the two due to built-in capabilities within the PubChem web interface (e.g., filtering function) that are not supplied through programmatic access. After converting the reference protocol into a prompt, we instruct GPT-4o to respond to the query or to generate the programmatic access. We compare the results against the gold and silver answers respectively.

Our paper is the first to evaluate the ability of GPT-4o in retrieving information from a biological database such as PubChem. The contributions of the paper are:

- Assessing the performance of GPT-4o in accessing the PubChem database on eight representative and commonly requested tasks [18], and qualitatively and quantitatively comparing the results against those obtained via the PubChem interface and through programmatic access.
- We show that retrieval through direct prompting is currently not satisfactory for all eight protocols, despite prompt enhancements, reflecting the lack of deep understanding of chemistry and limitation in access and retrieval modalities for the LLM.
- We show that code generation for the programmatic access is consistently more successful in retrieving the relevant data than prompting for direct retrieval when the programmatic access is available, and report success in 3 out of 5 cases.
- We outline current values and limitations of using a pre-trained general-purpose search-enabled LLMs in accessing databases and outline future directions to improve LLMs for database access.

## 2. Methods

### 2.1 The Task protocol and its adaptation into a prompt

To evaluate GPT-4o’s capabilities, we used eight PubChem retrieval protocols from reference [18]. Each of the tasks is outlined in Figures 1-8. A short description of the objective of each protocol appears at the top of each figure. For example, “Basic Protocol 1: Finding Genes and Proteins that Interact with a Given Compound”. We also added a description of the key steps involved in each search, e.g., 2D structure search, and filtering. The reference protocol steps for each task are outlined in the **upper left panel** of each figure. While the steps in the protocol are generic (applicable to any compound or any gene), each protocol is made specific per the details provided in reference [18]. For example, protocol #8 is applied to CID 60846, corresponding to valsartan.

**Figure 1.**
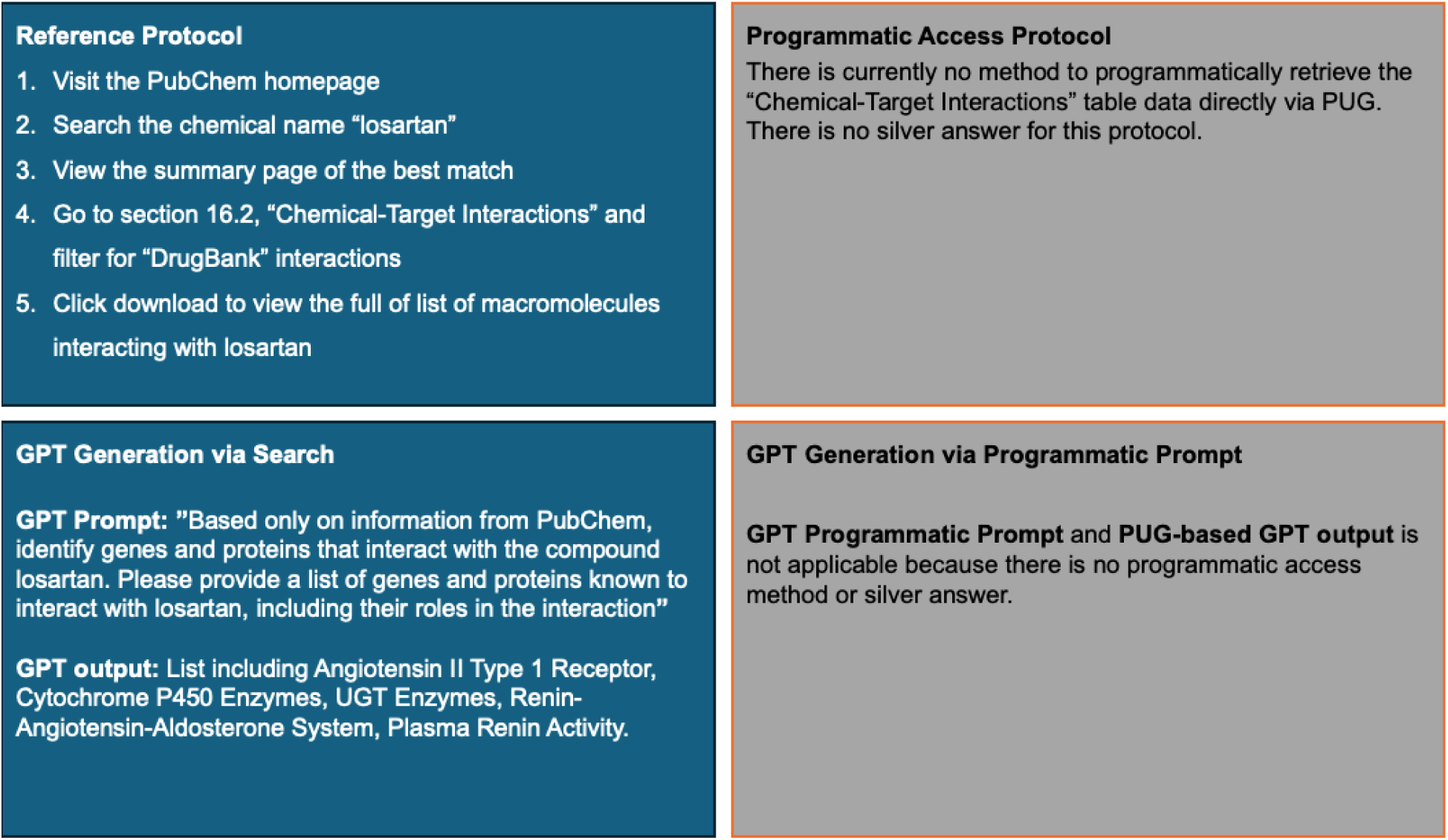
Protocol #1 for finding genes and proteins that interact with a given compound, made specific for losartan.

**Figure 2.**
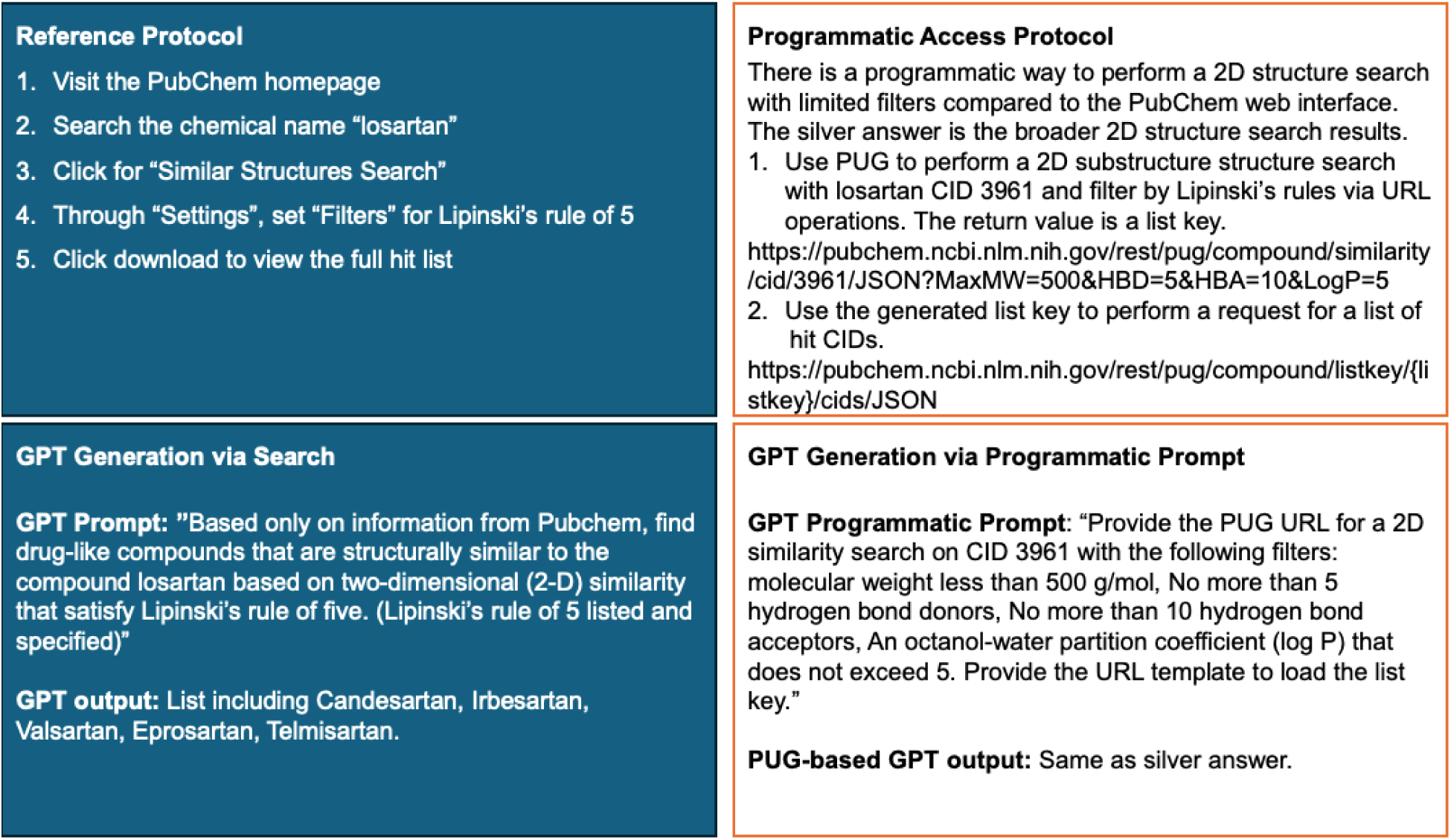
Protocol #2 for finding drug-like compounds similar to a query compound through 2-D similarity search, made specific for losartan filtered through Lipinski’s rule of 5.

**Figure 3.**
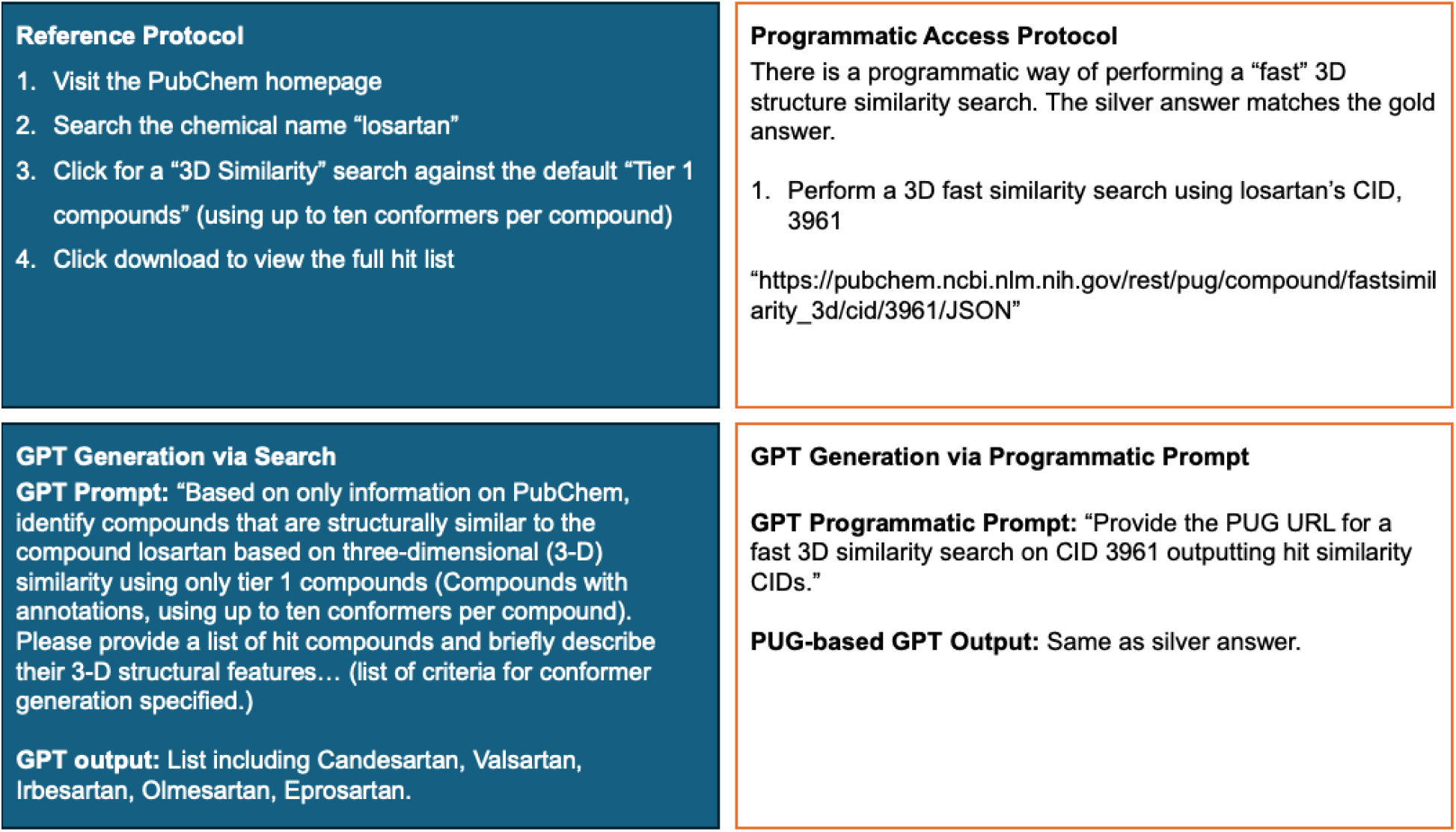
Protocol #3 for finding compounds similar to a query compound through 3-D similarity search, made specific for losartan.

**Figure 4.**
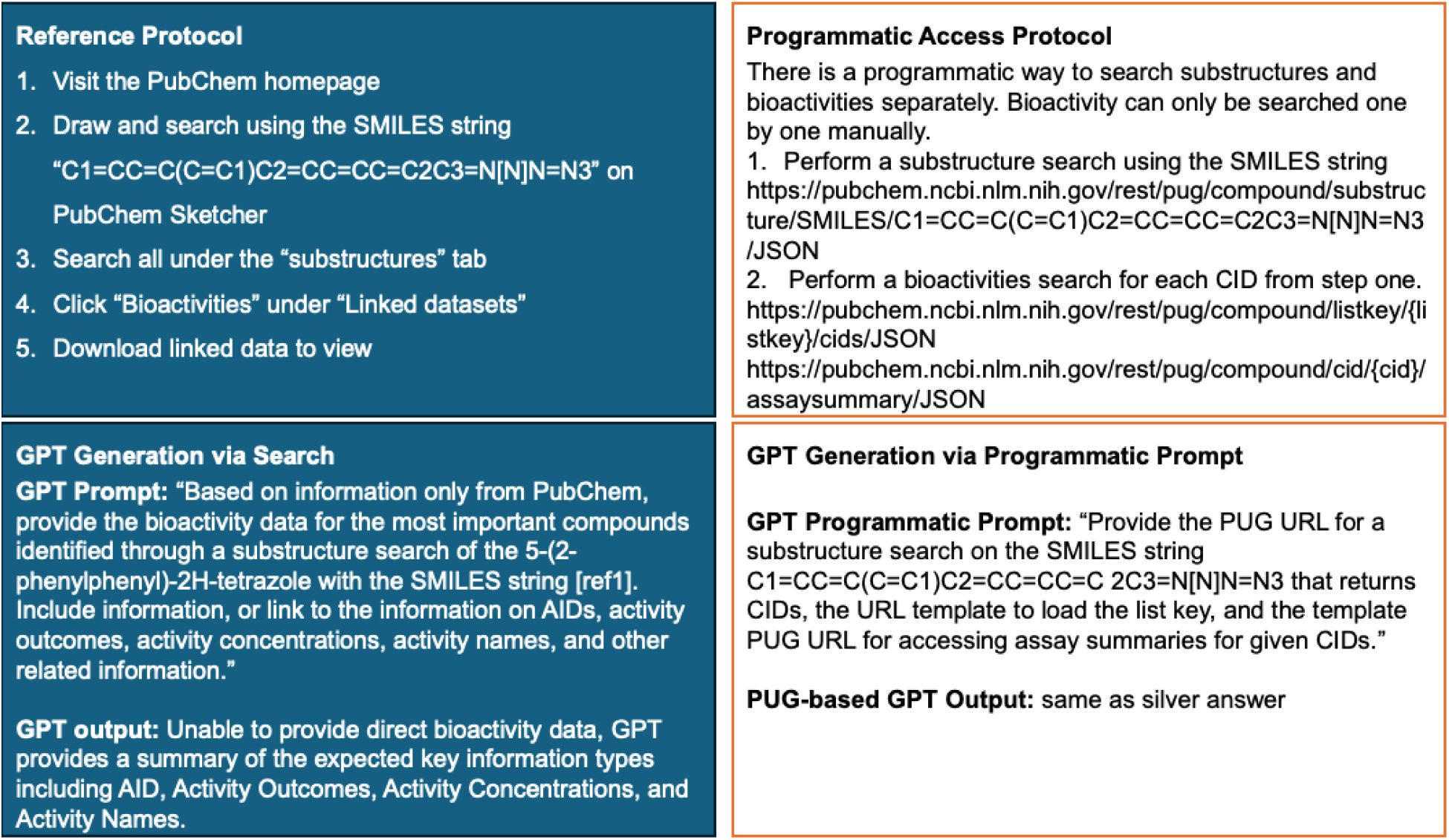
Protocol #4 for getting the bioactivity data for the hit compounds from substructure search, made specific for C1=CC=C(C=C1)C2=CC=CC=C2C3=N[N]N=N3.

**Figure 5.**
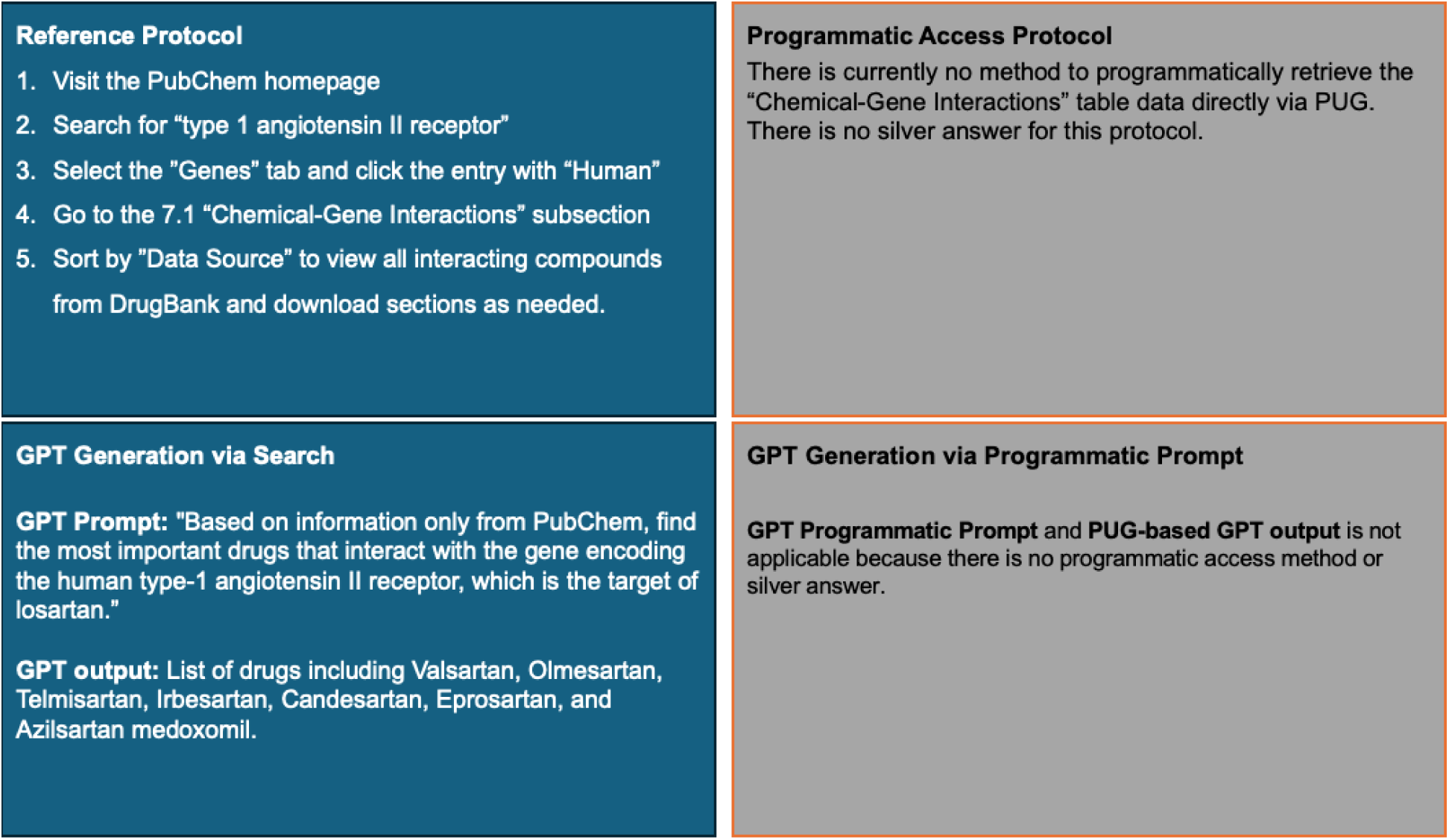
Protocol #5 for finding drugs that target a particular gene, made specific for the gene type 1 angiotensin receptor (human).

**Figure 6.**
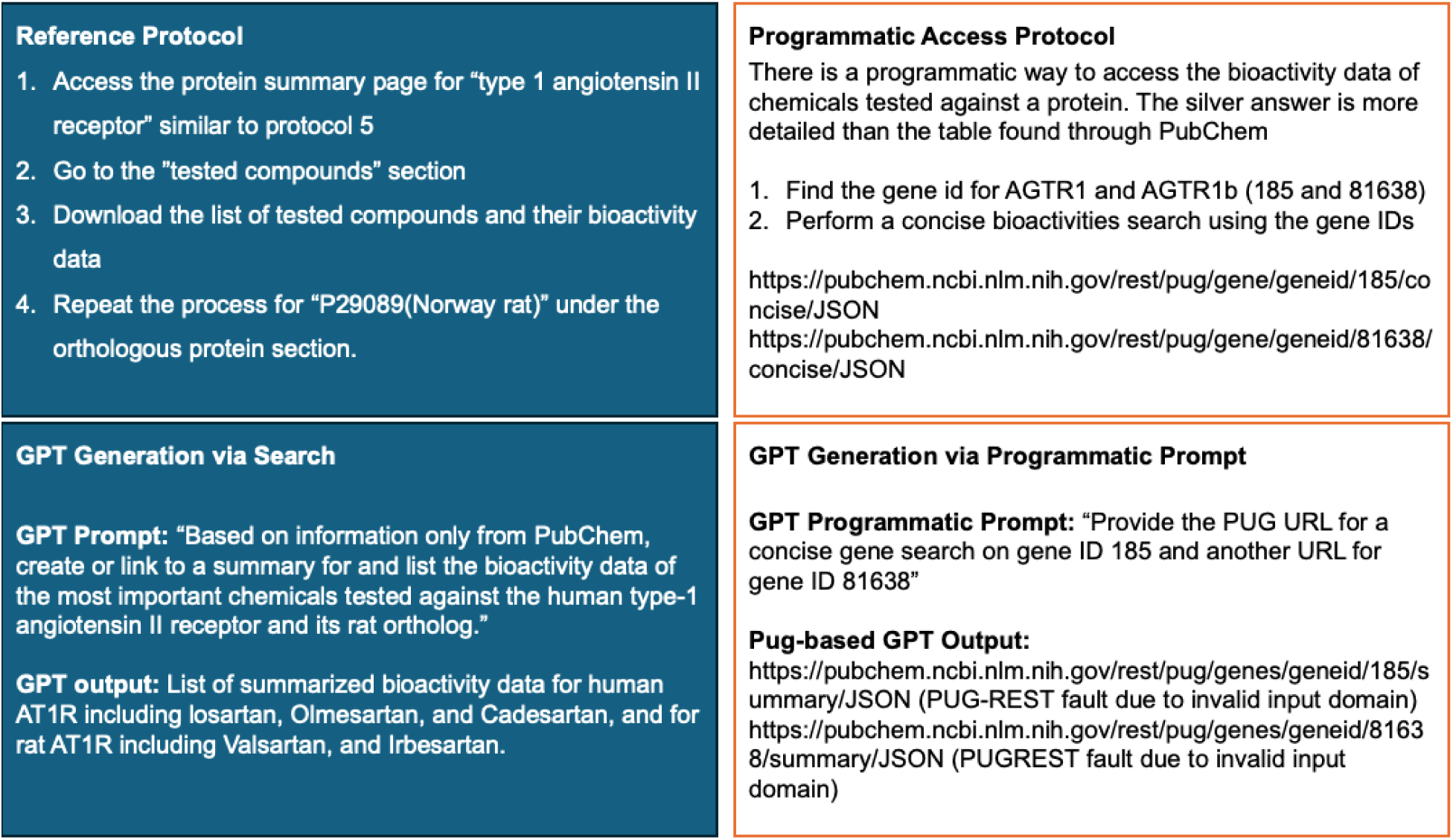
Protocol #6 for getting bioactivity data of all chemicals tested against a protein, made specific for the protein type 1 angiotensin II receptor (human and Norway rat).

**Figure 7.**
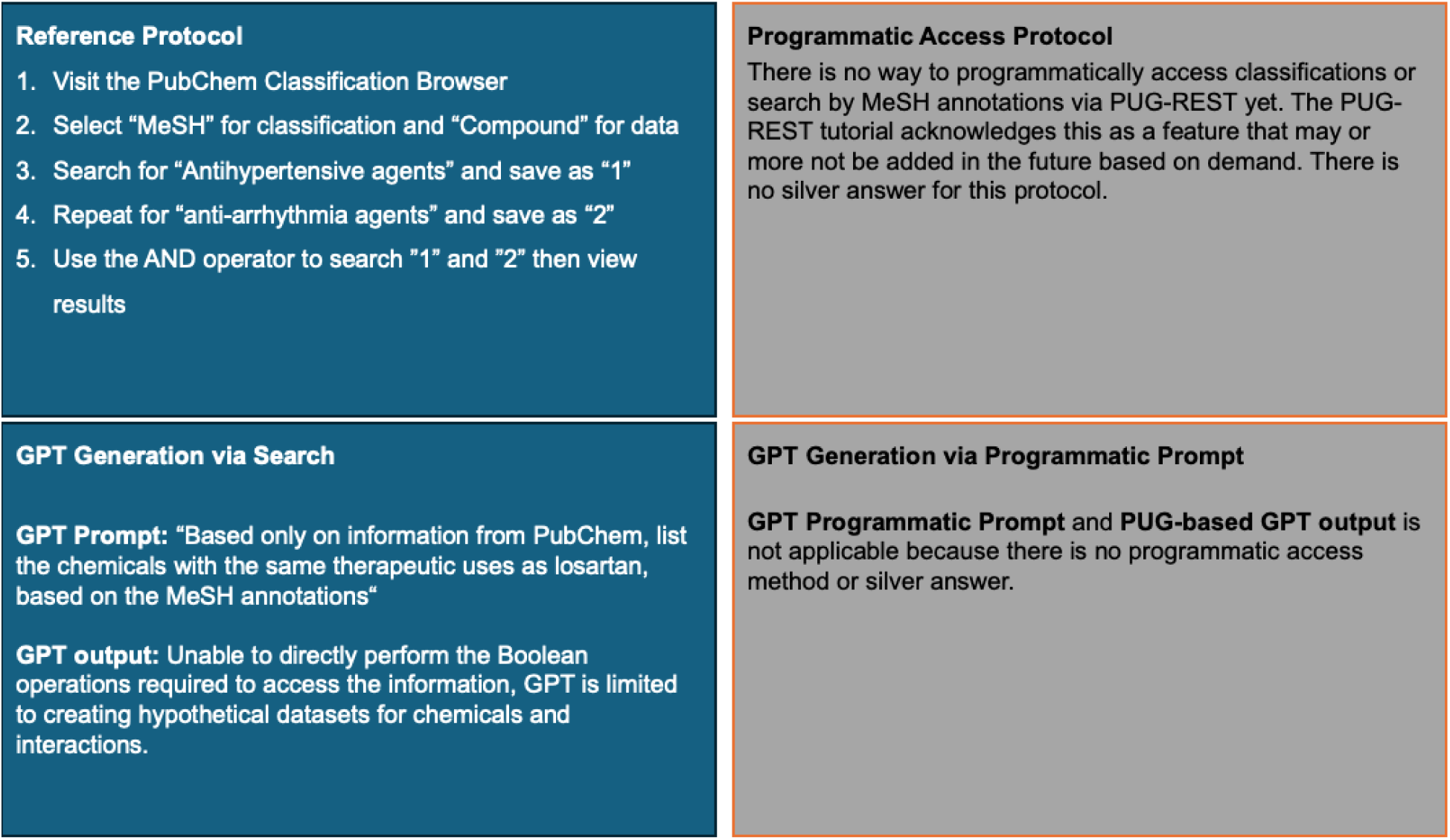
Protocol #7 for finding compounds annotated with classifications or ontological terms, made specific for “antihypertensive agents” and “anti-arrhythmia agents.”

**Figure 8.**
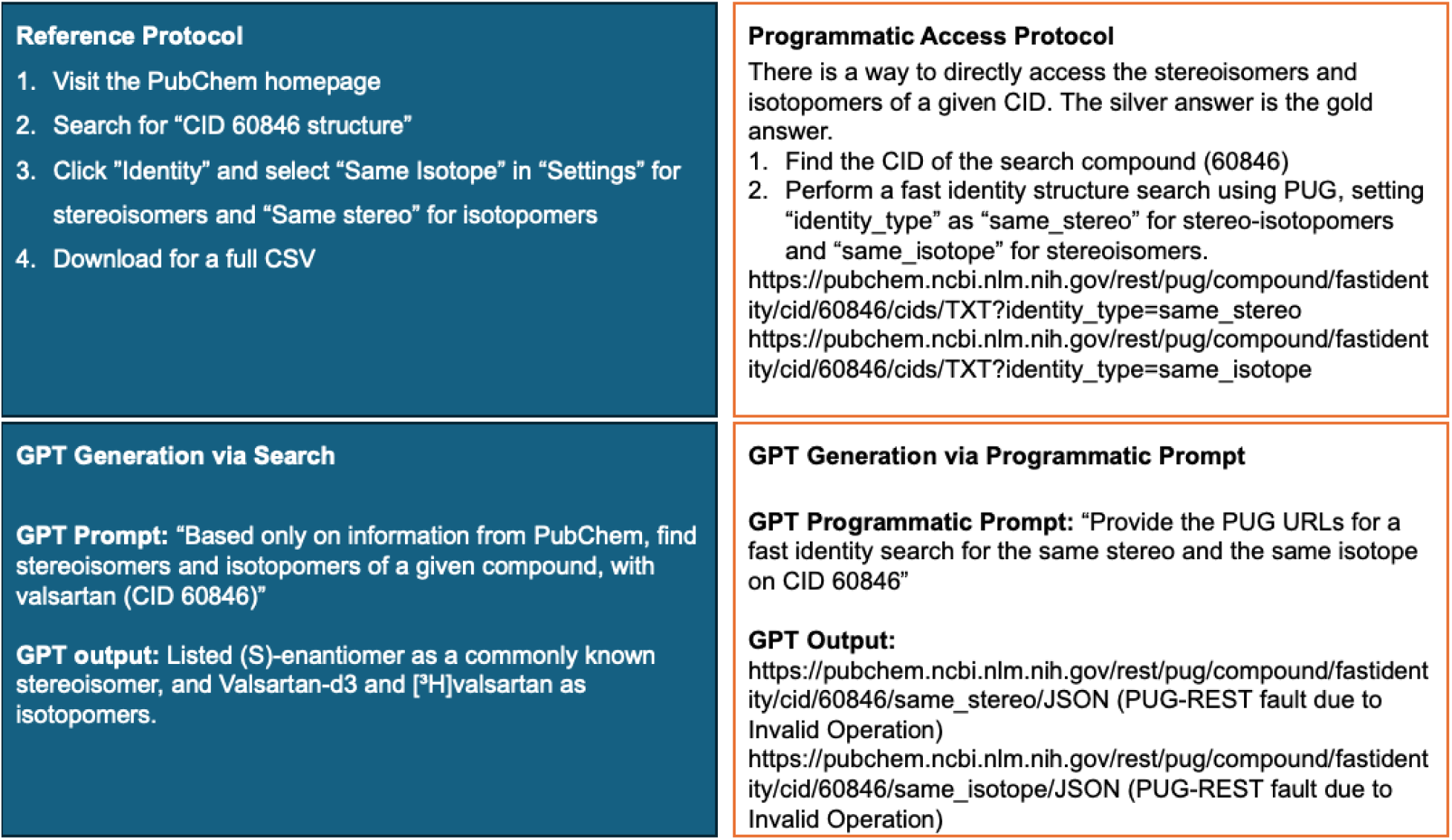
Protocol #8 for getting stereoisomers and isotopomers of a compound through identity search, made specific for a search on “CID 60846 structure.”

#### Initial Protocol Adaptation

Each protocol was formed into a prompt using the following process. The protocol’s specific details were utilized, and steps related to dynamic content handling via the PubChem web interface we ignored. Next, the question within the protocol was rephrased into a single, cohesive query for GPT-4o. To align with the referenced protocol, the prompts instructed GPT-4o to use only data from PubChem, explicitly avoiding external sources to maintain the accuracy and relevance of the retrieved data. In the cases where python code was generated in response to our first prompt, we modified the prompt to avoid any code generation. The lower left panel in Figures 1-8 displays the prompts used for each protocol and the summarized GPT-4o output.

#### Exploration of Enhanced Protocol

As GPT-4o is interactive, we examined feedback from the initial prompt, and judiciously prompt engineered the query by adding relevant context. Several types of enhancements were made. GPT-4o seemed sensitive to the question; we therefore refined our prompts to use certain phrases such as “list” vs “retrieve” to determine the phrasing that yielded the most accurate responses from GPT-4o. Another enhancement was through providing additional information. For instance, protocol #2 aims to find drug-like compounds similar to a query compound through 2-D similarity search, and then selects filtering, based on Lipinski’s rule of five, available through the PubChem interface to filter the results. In this case, we provided a detailed description for Lipinski’s rule. We further enhanced the prompt for example, by removing requests to computationally generate 2-D or 3-D models, a function available in PubChem but not through GPT-4o. When working with programmatic prompts, certain PubChem key terms such as “3-D structure search” in protocol 3, and “same stereo/isotope” in protocol #8 needed to be especially emphasized as URL operations, specifically “fast 3D similarity search” and “fast identity search” to generate the correct link. In the case of highly specific PUG operations, such as in protocol 5, the entire link template for the search from PubChem’s PUG-REST API is needed for a correct output. See Appendix for all enhanced prompts.

### 2.2 Protocol adaptation into a prompt with programmatic access

As PubChem provides programmatic access, we explore prompting GPT-4o to retrieve information via programmatic access and compare the retrieval results against our prompts described above that do not specifically instruct the GPT-4o to generate code. First, however, we manually explored PubChem’s PUG capabilities in servicing the protocol. This process allowed us to assess if programmatic access alone (not through the web interface) can secure the results attained through the reference protocols as we expected the PubChem web interface provided additional filtering capabilities not available through programmatic access. The upper right panels in Figures 1-8 describe the programmatic access available for each protocol. In some cases, such as in protocol #7, where the aim is to find compounds annotated with classifications or ontological terms, there is currently no programmatic access to classifications using the PUG-REST, per PubChem web documentation. Such a limitation prevents the GPT-4o from generating programmatic access to retrieve data. The bottom right of Figures 1-8 specify the prompt used on GPT-4o to generate URLs for programmatic access.

### 2.3 Selection of LLM model for evaluation

GPT-4o was selected for this study based on its performance across various metrics and user preferences evaluated on the Chatbot Arena [19]. This resource ranks LLM models based on human votes in categories such as overall performance, multi-turn conversations, handling longer queries, hard prompts, and coding capabilities. Users compare outputs from two anonymous models, side-by-side, and vote for the better response. GPT-4o ranks first in key categories including overall performance, hard prompts (overall and in English), responding to longer queries, and English proficiency. It ranks second to Claude in the multi-turn category, which is essential for maintaining coherent and contextually accurate conversations over multiple interactions, particularly for complex queries requiring step-by-step guidance. Additionally, GPT-4o’s ability to retrieve information from websites offers an advantage for accessing up-to-date data and supporting real-time information needs, which is critical for dynamic databases like PubChem. GPT-4o’s superior performance and user-preferred ranking makes it a suitable option for evaluating the retrieval tasks in this study.

### 2.4 Evaluation Method

With the execution of the basic protocol, the GPT-4o prompts (and their enhanced versions if utilized), the programmatic access, and the GPT-4o programmatic prompts, we can compare the results using the four methods. The execution of the basic protocol through the PubChem website per the instructions in reference [18] yields the “gold” answers to the queries. The execution of known programmatic access to execute the queries reflects the available answers through such modality, which we label as the “silver” answers. We therefore compare the GPT-4o responses to the gold answers when executing the GPT-4o prompts, and we also compare the results of running the GPT-4o programmatic prompts to the silver answers. As current GPTs cannot run the generated code, we manually evaluated the generated code.

## 3. Results and Discussion

Our study evaluates the responses of GPT-4o in response to prompting and code generation. We first describe our experiences with prompt engineering. Next, we provide a summary of our results, highlighting comparisons between the gold and silver answers, the gold answers and the prompts that did not specifically request using the programmatic access, and the silver answers and the prompts that specifically were designed to generate programmatic access. We then discuss the observed value of GPT-4o’s retrieval capabilities and outline a future vision for LLMs and chatbots for databases.

### 3.1 Prompt Engineering

Prompt engineering is now recognized as essential for obtaining accurate and relevant responses from LLMs [20].We found that minor changes to the wording of queries significantly impacts the model’s behavior. When querying GPT-4o to provide data from PubChem, using terms like “show” or “list” tends to yield better results than “analyze” or “retrieve,” which often lead to differing information or non-functional code. In some instances, as for protocol #3, 4, and 7, the prompt resulted in python code without the prompt requesting programmatic prompting. With enhanced prompting through interactive refinement, we added contextual information or definitions to clarify some protocols. For example, when enhancing the prompt for protocol 2, Lipinski’s rule of five was elaborated upon as “a molecular weight less than 500 g/mol, no more than 5 hydrogen bond donors, no more than 10 hydrogen bond acceptors, an octanol-water partition coefficient (log P) that does not exceed 5.”. Clarifications were also provided for “tier 1 compounds”, where the following was added to the prompt, “Generate a conformer model for each compound if it satisfies the following criteria: not too large (with ≤50 non-hydrogen atoms), not too flexible (with ≤15 rotatable bonds), has fewer than six undefined atom or bond stereocenters, has only a single covalently bonded unit (i.e., not a salt or a mixture), consists of only supported organic elements (H, C, N, O, F, Si, P, S, Cl, Br, and I), and contains only atom types recognized by the MMFF94s force field.” Importantly, adapting the reference protocol into one understandable by GPT-4o, and further enhancing it to yield better outputs or programmatic URLs is a task suited for all researchers, regardless of their coding proficiency.

### 3.2 Summary of Results for using GPT-4o on the protocols

Table 1 provides a summary of running the eight protocols using the four retrieval methods. When comparing the gold and silver answers, we were able to obtain the same answers in two cases, and partial overlaps in three cases, and no matches in three cases. The reason for partial or no match is due to additional capabilities available through the web interface that are not available through the programmatic access, or the lack of programmatic access.

**Table 1.** A summary of the results of evaluating GPT-4o on the eight protocols. Column 1 indicates the protocol number. Column 2 indicates if the gold answer matched the silver one, and column 3 provides a short explanation of why the two answers differed. Columns 4 and 5 summarize the results of the initial prompt or enhanced prompt, respectively. Columns 5 and 6 indicate if GPT-4o was able to create the URL for the programmatic access, and if the generated URL(s) results matched the silver answer.

|  | PubChem access without LLM |  | GPT-4o |  |  |  |
| --- | --- | --- | --- | --- | --- | --- |
|  | Web Interface vs Programmatic access |  | GPT-4o Results |  | Generation of URL for Programmatic access |  |
| Protocol | Gold matches silver answer? | If not, why? | Attained gold with initial prompt? | Attained gold with enhanced prompt? | Generated programmatic access URL? | Generated URL(s) are the silver answer? |
| 1 | No | No programmatic access method. | Partially, incomplete output with extraneous information | No improvement from initial prompt | N/A | N/A |
| 2 | Partially | Filtering through programmatic access yields a different result than filtering through the web interface | No, incorrect answers | No, shortened existing answer | Yes | Yes, but requires multiple steps |
| 3 | Yes |  | No, incorrect answers | No improvement from initial prompt | Yes | Yes |
| 4 | Partially | Programmatic access for bioactivity data is different than web access data | No, directed user to search on PubChem | No, created hypothetical data | Yes | Yes, but requires multiple steps |
| 5 | No | No programmatic access method. | Partially, 7 out of 14 drugs identified | No improvement from the initial prompt | No, faulty URL | N/A |
| 6 | Partially | Programmatic access is output is different from web output | Partially, unable to provide full answer | No improvement from the initial prompt | N/A | N/A |
| 7 | No | No programmatic access method | No, incorrect answers | No improvement from the initial prompt | N/A | N/A |
| 8 | Yes |  | No, incorrect answers | No improvement from the initial prompt | No, faulty URL | N/A |

When examining the output of GPT-4o due to our initial prompting, three cases resulted in partial matches to the gold answers, where only a subset of the results are provided. From examining the GPT-4o response, we noted different search sources were utilized other than PubChem, despite specific instructions to search from only PubChem. In the four other cases, the response was incorrect. In one case (protocol #4), GPT-4o directed the user to visit the PubChem website. Even with enhanced prompting, such as providing additional information or coaxing GPT-4o with different variations of the prompt, we did not witness material improvements in the responses. In one case (protocol #4), GPT-4o hallucinated by creating a fictitious table using randomly GPT-4o-generated numbers.

Examining the output of GPT-4o when instructed to generate the URL(s) for programmatic access, there was one case (protocol #7) where no programmatic access was possible. There were four correct URLs that matched the silver answer. In the remaining three cases, the generated URLs were faulty.

In summary, enhanced prompting did not yield improved results. However, quantitative analysis showed that such results are more useful to prompt follow-up interactions with the GPT-4o. Additionally, GPT-4o was better at generating the relevant URLs than performing search. Our analysis on the gold vs silver answers also revealed a gap in the capabilities of the programmatic access. Currently, GPT-4o is not sufficient to replace manual search on PubChem’s web interface.

### 3.3 Detailed results on the protocols

To enhance the understanding of the prior results, we describe in this section our experiences with GPT-4o in more detail, highlighting values and limitations that are specific to each case.

#### Protocol 1

The first protocol identifies genes and proteins that interact with losartan. Such data in PubChem originates from several sources including DrugBank [21], Drug-Gene Interaction database [22], and others. The PubChem web interface guides the user to go to section 16.2 (Chemical-Target Interaction), and filter against DrugBank interactions. The gold results (see full results in appendix) include several Cytochrome enzymes, UDP-glucuronosyltransferases, and others.

Based on prompting, GPT-4o provides correct yet incomplete results. The response is missing certain entries, e.g., Cytochrome P450 2C19, Cytochrome P450 2C8, and ATO-dependent translocase ABCB1. GPT-4o elaborates on each interaction with information external to PubChem and DrugBank such as Wikipedia (see partial results in Figure 1). The names of some interactions and proteins differ slightly from PubChem, indicating the results were acquired from external sources or training data. While the PubChem protocol had a strict definition of what qualifies as an interaction, and where the interactions must be sourced from, GPT-4o interpreted interactions as broader, and therefore included information about downstream effects of these interactions, listing “RAAS” and “Plasma Renin Activity” in the list of proteins and genes affected. Enhanced prompting for this protocol involved excluding downstream effects and protein or gene information and emphasizing the need for data to only be from PubChem. Unfortunately, any engineering to enhance the prompt led to incorrect answers with non-PubChem sources which strayed further from the established gold answer.

There is currently no method to retrieve the same information through programmatic access. Therefore, the silver answer is not available.

#### Protocol 2

The second protocol aims to find drug-like compounds similar to losartan based on a two-dimensional (2-D) similarity search using PubChem. The PubChem web interface guides users to perform a “Similar Structures Search” and apply Lipinski’s rule of 5 filters. The gold answer includes a detailed list of compounds that match the specified criteria (see full results in the appendix).

GPT-4o attempts to provide a list of compounds that are structurally similar to losartan while adhering to Lipinski’s rule of five. However, the results are incorrect when compared to the gold answer. None of GPT-4o’s outputs, such as Candesartan and Irbesartan, match the outputs from the gold answer. Looking into the external links provided by GPT-4o show that the model interpreted losartan as all types of losartan, including losartan Sodium. Enhanced prompting for this protocol involved specifying losartan as only “losartan” and not other variations of the drug. This enhanced prompt only shortened the output, leaving it with Candesartan, Irbesartan, and Valsartan, which were still incorrect compared to the gold standard.

To retrieve similar information programmatically, we used PUG to perform a 2-D substructure search for losartan with CID 3961 and applied filters for Lipinski’s rule of 5. The programmatic access involved using a URL to perform the search and obtain a list key, which was then used to request a detailed list of hit CIDs. This method provided comprehensive and accurate results that align with PubChem’s standards, but upon comparison with the gold answer, the retrieved information filtered on broader criteria, leading to a longer list of hit targets inclusive of the gold answer. This silver answer highlights a significant difference between searches done on PUG and the PubChem web interface.

We also prompted GPT-4o with a programmatic access approach to provide a PUG link for the 2-D similarity search. The prompt included specific criteria, such as molecular weight, hydrogen bond donors and acceptors, and partition coefficient limits according to Lipinski’s rule of five. GPT-4o generated the silver answer PUG URL. Although this is a multi-step protocol, requiring the user to manually input the generated list key for the final result, the programmatic prompt effectively provided the most relevant PUG URL templates for each step for detailed data access (refer to figure 2 for a full protocol summary).

#### Protocol 3

The third protocol aims to find compounds similar to losartan based on a three-dimensional (3-D) similarity search using PubChem. The data originates from PubChem’s chemical database. The PubChem web interface guides users to perform a “3D Similarity” search against the default “Tier 1 compounds” (compounds using up to ten conformers per compound). The gold standard results include a detailed list of compounds that match the specified criteria (see full results in the appendix).

GPT-4o attempts to provide a list of compounds that are structurally similar to losartan while adhering to the specified 3-D similarity criteria. However, the results are incorrect and contain discrepancies. The compounds listed by GPT-4o do not correspond to any of the 28 hit compounds from the gold answer. Additionally, Candesartan and Olmesartan listed by GPT-4o have incorrect CIDs when compared to PubChem. Attempting to enhance the prompt with details on the definition of “tier 1 compounds” (the default search filter applied by PubChem on similarity searches) does not change the correctness of the answer.

To retrieve similar information programmatically, we utilized PUG to perform a “fast” 3-D similarity search for losartan using CID 3961. The programmatic access involved using a URL to perform the search and obtain detailed compound data. This method provided comprehensive and accurate results that align with the gold answer.

We also prompted GPT-4o with a programmatic access approach to provide a PUG link for the fast 3-D similarity search. GPT-4o generated the correct PUG URL, which corresponds to the silver as well as gold answer. The programmatic prompt effectively provided the necessary link for data access (refer to figure 3 for a full protocol summary).

#### Protocol 4

The fourth protocol aims to get bioactivity data for hit compounds identified through a substructure search using a specific SMILES string in PubChem. The data originates from various sources within PubChem’s database. The PubChem web interface guides users to draw and search using the SMILES string “C1=CC=C(C=C1)C2=CC=CC=C2C3=N[N]N=N3” on the PubChem Sketcher, search under the “substructures” tab, click “Bioactivities” under “Linked datasets,” and download the linked data to view the bioactivity results (see full results in the appendix).

GPT-4o is limited in answering multimodal questions, such as those involving SMILES formulas and drawn structures. When GPT-4o is given the SMILES formula, it cannot directly perform a structure search on PubChem. Instead, it instructs users on how to conduct the search and provides limited information through external links. As a result, GPT-4o can only provide a summary of the expected key information types, including AID, Activity Outcomes, Activity Concentrations, and Activity Names, but not the direct bioactivity data itself. Attempting to enhance the prompt led to code generation which created hypothetical data unrelated to actual PubChem data.

To retrieve similar information programmatically, we utilized PUG to perform a substructure search using the given SMILES string. This was followed by performing a search for bioactivities for each CID from the initial search results. Alternatively, a CSV file of all bioactivities of CIDs can be downloaded if all CIDs are listed in the same link. This method involved using specific URLs to first obtain a list of CIDs from the substructure search and then retrieve detailed bioactivity data for each CID. This approach is labor-intensive and more difficult to interpret than the web interface, but provides highly comprehensive results that align with PubChem.

We also prompted GPT-4o with a programmatic access approach to provide a PUG link for the substructure search using the given SMILES string. The prompt included specific criteria to return CIDs, load the list key, and access assay summaries for the given CIDs. GPT-4o generated the correct PUG URLs, allowing for accurate retrieval of compound and bioactivity data, corresponding to the silver answer (refer to figure 4 for a full protocol summary).

#### Protocol 5

The fifth protocol aims to find drugs that target the human type-1 angiotensin II receptor gene. The data originates from several sources within PubChem, including DrugBank. The PubChem web interface guides users to search for “type 1 angiotensin II receptor,” select the “Genes” tab and click the entry with “Human,” navigate to the 7.1 “Chemical-Gene Interactions” subsection, sort by “Data Source” to view all interacting compounds from DrugBank, and download the necessary sections (see full results in the appendix).

GPT-4o provides a list of drugs that interact with the gene encoding the human type-1 angiotensin II receptor. The response is correct but incomplete. GPT-4o lists the drugs Valsartan, Olmesartan, Telmisartan, Irbesartan, Candesartan, and Azilsartan medoxomil, which is only 7 out of the 14 total drug interactions from DrugBank. Attempting to enhance the prompt by asking for a complete list, or all of the data lead to faulty code generation or an output unchanged from the initial output.

There is no programmatic way to directly access the gene-chemical interactions table. There is no silver response.

#### Protocol 6

The sixth protocol aims to get the bioactivity data of all chemicals tested against the human type-1 angiotensin II receptor (AT1R) and its rat ortholog. The data originates from various sources within PubChem. The PubChem web interface guides users to access the protein summary page for “type 1 angiotensin II receptor,” navigate to the “tested compounds” section, download the list of tested compounds and their bioactivity data, and repeat the process for “P29089 (Norway rat)” under the orthologous protein section (see full results in the appendix).

GPT-4o provides a summary of the bioactivity data for the most important chemicals tested against the human type-1 angiotensin II receptor and its rat ortholog. GPT-4o cannot directly search PubChem and thus only provides possible bioactivity types, a general overview of the AT1R receptor, and listings of human AT1R interactions for compounds losartan, Olmesartan, and Candesartan, and rat AT1R interactions for compounds Valsartan and Irbesartan. GPT-4o does not explain how it shortlisted these compounds, and descriptions of the compounds are generic and do not reflect the gold answer. Enhanced prompting by re-querying and asking for exact lists of compounds did not improve the quality of the responses for this protocol.

To retrieve similar information programmatically, we utilized PUG to perform a concise bioactivities search using the gene IDs 185 and 81638, corresponding to AGTR1 and AGTR1b respectively. This method involved finding the relevant gene IDs and using URLs to obtain detailed bioactivity data.

We also prompted GPT-4o with a programmatic access approach to provide PUG links for concise gene searches on gene ID 185 and gene ID 81638. The prompt included specific criteria to return a summary of bioactivity data for these gene IDs. GPT-4o generated incorrect PUG URLs due to an invalid input domain (refer to figure 6 for a full protocol summary).

#### Protocol 7

The seventh protocol aims to find compounds annotated with specific classifications or ontological terms using PubChem. The data originates from PubChem’s chemical database. The PubChem web interface guides users to the Classification Browser, selecting “MeSH” for classification and “Compound” for data, searching for “Antihypertensive agents” and saving as “1,” repeating for “anti-arrhythmia agents” and saving as “2,” and using the AND operator to search for results matching both classifications (see full results in the appendix).

GPT-4o cannot perform a direct search on PubChem and is thus unable to use the Boolean search function required for this protocol. The original protocol requires the user to create two separate searches and use the Boolean operator AND to find results that match both statements. While PubChem has 68 results for this search, none of them correspond to GPT-4o’s answers. GPT-4o instead generates hypothetical datasets of chemicals and interactions, which do not accurately reflect the actual data available in PubChem. Enhanced prompting did not improve the quality of the responses for this protocol.

There is currently no way to programmatically access classifications or search by MeSH annotations via PUG-REST. The PUG-REST tutorial acknowledges this as a feature that may be added in the future based on demand. Thus, there is no silver answer for this protocol through programmatic access.

#### Protocol 8

The eighth protocol aims to find stereoisomers and isotopomers of a compound using an identity search in PubChem. The data originates from PubChem’s comprehensive chemical database. The PubChem web interface guides users to search for “CID 60846 structure,” select “Identity” and select “Same Isotope” in “Settings’’ for stereoisomers and “Same stereo” for stereoisotopomers and download the results as a CSV file (see full results in the appendix).

GPT-4o provides a list of stereoisomers and isotopomers for Valsartan (CID 60846). The response includes the (S)-enantiomer as a commonly known stereoisomer, and Valsartan-d3 and [3H]Valsartan as isotopomers. However, the names and formatting of these chemical compounds differ from PubChem’s data, indicating discrepancies. The response does not match the gold answer from PubChem. Attempting to enhance the prompt by changing “find” to “list” or “retrieve” led to faulty code generation. Re-querying for the correct answer leads to a change in output compounds that was still incorrect.

To retrieve similar information programmatically, we used PUG to perform a fast identity structure search for stereoisomers and isotopomers using the CID 60846. This method involved setting “identity_type” as “same_stereo” for stereoisotopomers and “same_isotope” for stereoisomers and using specific URLs to obtain the silver answer which matched the gold answer.

We prompted GPT-4o with a programmatic access approach to provide PUG links for a fast identity search using the same stereo and same isotope settings for CID 60846. However, the output generated by GPT-4o was incorrect, resulting in PUG-REST faults due to invalid operations. (refer to figure 8 for a full protocol summary)

### 3.4 Observed Limitations of GPT-based retrieval

In evaluating GPT-4o on the eight protocols, we observed several limitations that contributed to the difficulty of using GPT-4o for retrieval and in realizing the gold and silver answers. One limitation is that GPT-4o was not able to follow instructions. For example, it did not necessarily limit its search to PubChem. GPT-4o searched and cited external sources such as Wikipedia to support its answers. For example, this issue is present in protocols 1, 5, and 6, where Wikipedia is directly cited. Further, the model could not effectively apply filters or rules, such as the rule of five or Boolean operations, demonstrating limitations in following complex, conditional instructions. Even with enhanced prompting (e.g., specifying Lipinksi’s rule of 5), GPT-4o could not apply such knowledge effectively, indicating the lack of fundamental understanding of chemistry or utilizing this knowledge to effectively filter the results.

Another limitation is the lack of multi-modal capabilities within GPT-4o. Search cannot be performed using drawings of compounds, and we did not observe positive results when using SMILES strings. Another limitation is GPT-4o’s lack of direct access to code bases in modules such as PubChemPy and RDKit. Such modules can aid in filtering and aggregation, capabilities currently built into the PubChem web interface. Even with code generation (not execution), the generated URLs were not always generated correctly. Yet, another limitation is the variability in responses due to the input prompt. For example, we observed sensitivity to keywords in the input prompt. When providing partial answers, such as protocols 1, 5 and 6, GPT-4o arbitrarily selects subsets of results as relevant without clear justification for excluding others. This inconsistency poses a challenge for reproducibility and reliability and potentially reflects the sampling aspect of the LLMs.

### 3.5 Observed value of GPT-based retrieval

Despite its tremendous shortcomings in generating the gold and silver answers, there were key observed values in utilizing GPT-4o. GPT-4o shows promise in performing semantic searches and presents an advantage over traditional search engines such as Google, especially for queries requiring nuanced understanding or involving multiple search steps. For example, searching protocol #1 in Google results in hits to multiple pages, including the PubChem page for the relevant compound, published paper, DrugBank, the Rat Genome Database, and others. With the PubChem page hit, a user must visit the various page sections to identify the relevant information. Another perceived advantage is the interactivity between human and GPT while retaining the context and history of a given conversation. Coherent and continuous dialogues enable users to build on previous queries without restarting. This personalized approach is beneficial for complex queries that require step-by-step guidance and contextual understanding. (in search and programmatic access). When offering guidance or step-by-step instructions, GPT-4o helps users in understanding of the processes involved in retrieving and analyzing chemical information. For example, in protocol 4, where GPT-4o could not directly perform a substructure search using a SMILES string, it instead guided users on how to conduct the search manually through the web interface. This instructional capability enhances user comprehension of PubChem’s functions and encourages users to perform complex searches independently.

When prompting GPT-4o for the URLs for programmatic access, the provided links resulted in correct answers in three of the five cases where programmatic access was available. An important outcome is that the programmatic link leads to only PubChem-based information. GPT-4o’s ability to generate partial code promises to enhance user productivity, particularly for a coder unfamiliar with the PubChem programmatic access. The generated code, even if requiring refinement and debugging, provides a foundation that reduces the effort required to set up the basic framework. For example, in protocol 2, GPT-4o provided python code for implementing Lipinski’s rule of five. This capability not only accelerates the development process but also allows users to focus more on refining and analyzing data rather than on preliminary coding tasks.

### 3.6 Future vision for LLMs for PubChem and other databases

There are two potential parallel future directions. One direction is to enhance existing general-purpose pre-trained LLMs for database access through various improvements. Fundamentally, intelligently using a general-purpose LLM for accessing biological and chemical databases requires deeper, nauseated understanding of chemistry, biology and biochemistry, and learning the application of these disciplines to the retrieved results. Enhancing different search modalities will enable handling queries involving SMILES notation and structural drawings, as encountered in protocol 4. Providing the LLM with access to external analytical tools such as RDkit and PubChemPy would guarantee the LLM to carry out searches through the PubChem database rather than external sources. Integrating knowledge graphs that powers a database such as PubChem by linking chemical compounds to their properties, interactions, and literature would prevent misinterpretations of key words and operations. Enhancing semantic search capabilities would improve an LLM’s ability to interact with users, handle complex queries and retrieve data. Designing a system for task decomposition would allow LLMs to break down multi-step queries into manageable sub-tasks, which can then be picked up by humans if any steps go wrong. Personalization for individual users based on experience level and research topic is unique to LLM memory and can provide more tailored and relevant information. In combination with validation mechanisms, users can be sure the retrieved data meets specific criteria even for highly technical searches.

Another possibility is to create chatbots specialized for a specific discipline (biochemist expert!) or for specific database(s), e.g., a chatbot for PubChem, or a combination thereof. Using a specialized LLM for interactive PubChem search combines the ease of communicating in natural language with the reliable and exact search capabilities of PubChem. Using a specialized chatbot as a co-pilot in instructing users on how to perform search or providing general links to relevant information on PubChem improves overall accessibility to the database’s resources. These advancements will collectively enhance the functionality and user experience of LLMs in scientific research. Enriching such an LLM with additional biological, chemical, or biochemical knowledge can even yield improved performance.

## Conclusion

We are currently in the early stages of exploring the use of LLMs as an integral part of science and discovery. Here, the study evaluated the capabilities and limitations of a pre-trained, searched-enabled LLM in accessing and retrieving data from the PubChem database. The results in Table 1 indicate limitations that prevent current LLMs from being sufficient as a standalone PubChem search and retrieval tool. Even with enhanced prompting, it was clear that the GPT-4o lacked the domain knowledge to interpret and filter the results. While GPT-4o was successful in generating some of the URLs for programmatic access, it was not successful in all cases. Our perceived value is that the GPT-4o can fulfill the role of a (sometimes trusted) tutor. This initial study highlights the need for significantly improved LLMs with domain expertise, combined with LLMs specialized in database retrieval. We suspect that the latter agents must understand not just the semantics of the data stored within the database, but the intricate relationships among its entities (e.g., the underlying knowledge graph). In addition, we expect that improvements in general-purpose LLMs, such as instruction following, multi-modal capabilities, and others can pave the way to smarter chatbots for database access. However, as the technology is moving swiftly, perhaps we will be pleasantly surprised within the next decade to have one intelligent agent capable of it all!

## ASSOCIATED CONTENT

Supporting File 1.pdf: This file contains snippets of the gold answers, and GPT responses for the 8 protocols.

Supporting File 2.xlsx: This file details the gold answers for each of the protocols

## AUTHOR INFORMATION

### Author Contributions

SH conceived the study and AS provided the implementation. Both authors analyzed the results. The manuscript was written through contributions of all authors. All authors have given approval to the final version of the manuscript.

### Data and Software Availability

The gold and silver answers were acquired through searches on PubChem, utilizing protocols specified in “Exploring Chemical Information in PubChem” by Sunghwan Kim. These protocols were modified for comprehension by GPT-4 (ChatGPT-4o). The outputs generated by GPT-4o were obtained using ChatGPT’s GPT-4o model.

The data and protocols used in this study are accessible as follows:

- PubChem Data: The raw data and search results can be accessed via the PubChem database. This data is publicly available and can be freely accessed.
- Modified Protocols: The modified protocols for querying PubChem using GPT-4o are essential for the implementation of our methods. While exact reproduction of the results generated by GPT-4o may vary due to the inherent variability of the model, the steps outlined in the appendix provide a comprehensive guide to closely replicate our approach. These steps include detailed instructions and examples that can be followed to achieve similar outcomes, recognizing that some variability in the responses is expected.
- GPT-4o Outputs: The outputs generated by GPT-4 (ChatGPT-4o) are proprietary and can be obtained following the steps in the appendix using the ChatGPT platform provided by OpenAI.

The GPT-4o outputs are proprietary because it uses a commercial platform (ChatGPT) provided by OpenAI. The outputs generated by GPT-4o use proprietary algorithms and models developed by OpenAI. These outputs are subject to OpenAI’s terms of service and licensing agreements, which restricts the public sharing of the raw output data.

For review purposes, the ChatGPT platform can be accessed through OpenAI’s ChatGPT website. The detailed protocols and data sources can be found in the appendix and ensure aspects of the study can be replicated.

## ACKNOWLEDGMENT

The authors are grateful for feedback from the members of the Hassoun Lab on an initial version of the paper, and for the funding form the Army Research Office, MURI program.

