## Supporting File 1 for "Evaluation of search-enabled Pre-trained Large Language Models on retrieval tasks for the PubChem Database"

Ash Sze and Soha Hassoun
Tufts University

Supporting File 1

This file contains snippets of the gold answer and the GPT response for each of the eight protocols. The GPT prompt is listed in Figures 1-8. For reference, the complete gold answers are provided in Supporting File 2.

### Protocol 1: Finding Genes and Proteins that Interact with a Given Compound

#### Snippet of Gold Answer

| id | cid | dsn | cmpdname | srccmpd | srccmpdname |
|---|---|---|---|---|---|
| 446239 | 3961 | DrugBank | Losartan | DB00678 | Losartan |
| 446240 | 3961 | DrugBank | Losartan | DB00678 | Losartan |
| 446436 | 3961 | DrugBank | Losartan | DB00678 | Losartan |
| 446437 | 3961 | DrugBank | Losartan | DB00678 | Losartan |
| 446438 | 3961 | DrugBank | Losartan | DB00678 | Losartan |
| 446439 | 3961 | DrugBank | Losartan | DB00678 | Losartan |
| 446440 | 3961 | DrugBank | Losartan | DB00678 | Losartan |
| 446441 | 3961 | DrugBank | Losartan | DB00678 | Losartan |
| 446442 | 3961 | DrugBank | Losartan | DB00678 | Losartan |

#### GPT Generation (Bottom-Left of Figures 1-8)

“Based on information from PubChem and DrugBank, Losartan interacts with several genes and proteins.

Here are the key interactions:

1. **Angiotensin II Type 1 Receptor (AT1R)**: Losartan acts primarily by blocking the AT1 receptor, which prevents angiotensin II from binding and causing vasoconstriction. This action helps lower blood pressure and reduce the workload on the heart.
2. **Cytochrome P450 Enzymes (CYP2C9, CYP3A4)**: Losartan is metabolized in the liver by cytochrome P450 enzymes, particularly CYP2C9 and CYP3A4. This metabolism is crucial for converting Losartan into its active metabolite, E-3174, which has a more potent effect on blocking the AT1 receptor.
3. **UGT Enzymes (UGT1A1, UGT1A3, UGT1A10, UGT2B7, UGT2B17)**: These UDP-glucuronosyltransferase (UGT) enzymes are involved in the glucuronidation of Losartan, facilitating its excretion from the body.
4. **Renin-Angiotensin-Aldosterone System (RAAS)**: Losartan impacts this system by inhibiting the effects of angiotensin II, leading to decreased aldosterone secretion, reduced sodium reabsorption, and ultimately lower blood pressure.
5. **Plasma Renin Activity**: By blocking the feedback mechanism of angiotensin II, Losartan indirectly increases plasma renin activity. This increase is a compensatory response to the blockade of the RAAS pathway.

These interactions highlight the multi-faceted role of Losartan in treating hypertension and related cardiovascular conditions by targeting specific receptors and enzymes in the body (DrugBank) (DrugBank) (Wikipedia).”

### Protocol 2: Finding Drug-Like Compounds Similar to a Query Compound through a Two-Dimensional (2-D) Similarity Search

**Snippet of Gold Answer**

| cid | cmpdname | cmpdsynonym | mw | mf | polararea | complexity | xlogp |
|---|---|---|---|---|---|---|---|
| 3961 | Losartan | losartan\|114798-26- | 422.9 | C22H23ClN6O | 92.5 | 520 | 4.3 |
| 108185 | Losartan carboxylic | Losartan carboxylic | 436.9 | C22H21ClN6O2 | 110 | 590 | 5 |
| 9823584 | 4-Chloro-2-propyl-1-[ | SCHEMBL855\|CHEM | 422.9 | C21H19ClN6O2 | 110 | 576 | 4.5 |
| 9844369 | [5-chloro-2-propyl-3-[ | SCHEMBL962\|L001 | 408.9 | C21H21ClN6O | 92.5 | 506 | 3.7 |
| 9846177 | [2-propyl-3-[[4-[2-(2H | CHEMBL171524\|SC | 442.4 | C22H21F3N6O | 92.5 | 589 | 3.7 |
| 9867801 | Losartan metabolite | 141675-57-2\|Losarta | 438.9 | C22H23ClN6O2 | 113 | 552 | 3 |
| 9911047 | Losartan metabolite | 141675-59-4\|Losarta | 438.9 | C22H23ClN6O2 | 113 | 552 | 3 |
| 9979425 | [5-butyl-2-(hydroxymethyl)-1-[[4-[2-(2H-tetra | | 418.5 | C23H26N6O2 | 113 | 532 | 2.5 |
| 10341959 | [5-butyl-2-(hydroxymethyl)-3-[[4-[2-(2H-tetra | | 418.5 | C23H26N6O2 | 113 | 532 | 2.5 |
| 11270805 | [2-butyl-5-chloro-3-[4 | SCHEMBL2927903 | 408.9 | C21H21ClN6O | 92.5 | 506 | 4.4 |
| 11531866 | [2-[(E)-but-1-enyl]-5-c | SCHEMBL683439\|S | 420.9 | C22H21ClN6O | 92.5 | 558 | 4 |
| 11598393 | [5-(1,1,2,2,2-pentaflu | CHEMBL170944\|SC | 492.4 | C23H21F5N6O | 92.5 | 677 | 4.4 |
| 14962867 | [5-(2,5-dimethylpyrro | CHEMBL307432\|BD | 467.6 | C27H29N7O | 97.4 | 649 | 4.3 |

**GPT Response**

"Based on information from PubChem and other databases, several drug-like compounds structurally similar to Losartan satisfy Lipinski's rule of five. These compounds have a molecular weight of less than 500 g/mol, no more than 5 hydrogen bond donors, no more than 10 hydrogen bond acceptors, and an octanol-water partition coefficient (log P) that does not exceed 5. Here is a list of such compounds along with their molecular properties:

1. **Candesartan**
    - Molecular Weight: 440.46 g/mol
    - Heavy Atom Count: 32
    - Rotatable Bond Count: 6
    - Hydrogen Bond Donor Count: 2
    - Hydrogen Bond Acceptor Count: 8
    - Polar Surface Area: 126.5 Å²
    - Molecular Complexity: 676
    - XLogP: 3.7
2. **Irbesartan**
    - Molecular Weight: 428.53 g/mol

- Heavy Atom Count: 31
- Rotatable Bond Count: 6
- Hydrogen Bond Donor Count: 1
- Hydrogen Bond Acceptor Count: 6
- Polar Surface Area: 95.9 Å²
- Molecular Complexity: 579
- XLogP: 4.1

3. **Valsartan**
    - Molecular Weight: 435.5 g/mol
    - Heavy Atom Count: 30
    - Rotatable Bond Count: 8
    - Hydrogen Bond Donor Count: 2
    - Hydrogen Bond Acceptor Count: 5
    - Polar Surface Area: 111 Å²
    - Molecular Complexity: 602
    - XLogP: 3.9
4. **Eprosartan**
    - Molecular Weight: 424.5 g/mol
    - Heavy Atom Count: 30
    - Rotatable Bond Count: 8
    - Hydrogen Bond Donor Count: 1
    - Hydrogen Bond Acceptor Count: 7
    - Polar Surface Area: 104.7 Å²
    - Molecular Complexity: 557
    - XLogP: 4.5
5. **Telmisartan**

- Molecular Weight: 514.6 g/mol (slightly exceeds the 500 g/mol rule)
- Heavy Atom Count: 38
- Rotatable Bond Count: 6
- Hydrogen Bond Donor Count: 1
- Hydrogen Bond Acceptor Count: 6
- Polar Surface Area: 83.9 Å²
- Molecular Complexity: 668
- XLogP: 7.0 (exceeds the log P rule)

While Telmisartan does not strictly satisfy the rule of five due to its molecular weight and log P value, it is structurally similar to Losartan and included for comparative purposes.

These compounds are selected based on their structural similarity to Losartan and compliance with most of the criteria of Lipinski's rule of five. Each compound has unique properties that might be beneficial for different therapeutic applications, particularly in managing hypertension and related cardiovascular conditions (DrugBank) (DrugBank) (ChemSpider)."

### Protocol 3: Finding Compounds Similar to a Query Compound through a Three-Dimensional (3-D) Similarity Search

**Snippet of Gold Answer**

| cid | cmpdname | cmpdsynonym | mw | mf | polararea | complexity | xlogp |
|---|---|---|---|---|---|---|---|
| 3961 | Losartan | losartan\|114798-26-4 | 422.9 | C22H23ClN6O | 92.5 | 520 | 4.3 |
| 10597249 | [1-butyl-3-methyl-5-[[ | CHEMBL440358\|BDI | 402.5 | C23H26N6O | 92.5 | 514 | 3.7 |
| 19815401 | 2-butyl-5-hydroxymet | CHEMBL315094\|2-b | 456.5 | C23H23F3N6O | 92.5 | 603 | 4.2 |
| 15698108 | 5-[2-[4-[[2-butyl-4-chl | CHEMBL86207\|SCH | 446.9 | C24H23ClN6O | 103 | 635 | 4.5 |
| 130762 | 4'-((2-Butyl-4-chloro-5 | Exp7711\|114798-27- | 398.9 | C22H23ClN2O3 | 75.4 | 498 | 4.6 |
| 10453569 | 3-[4-[[2-Butyl-4-chlor | CHEMBL129225\|BDI | 399.9 | C21H22ClN3O3 | 88.2 | 501 | 3.9 |
| 10363575 | 2-[4-[[2-Butyl-4-chlor | CHEMBL129170\|BDI | 399.9 | C21H22ClN3O3 | 88.2 | 501 | 3.6 |
| 10001012 | 3-[4-[[2-Butyl-4-chlor | CHEMBL340682\|BDI | 399.9 | C21H22ClN3O3 | 88.2 | 501 | 3.5 |
| 11756211 | [2-butyl-5-iodo-3-[[4-[ | CHEMBL313715\|BDI | 514.4 | C22H23IN6O | 92.5 | 520 | 4 |

**GPT Generation**

“To identify compounds structurally similar to Losartan using PubChem's 3-D similarity search, we utilize the "Similar Conformers" approach. This method relies on two main 3-D similarity metrics: the shape Tanimoto (ST) and color Tanimoto (CT) scores. Compounds are considered similar if they have an ST score of 0.8 or greater and a CT score of 0.5 or greater.

Based on the tier 1 criteria (compounds with annotations, using up to ten conformers per compound), here are some hit compounds structurally similar to Losartan:

1. **Candesartan (CID 10844100)**
   - **3-D Structural Features**: Candesartan shares a biphenyl tetrazole structure similar to Losartan, with additional ester and carboxyl groups enhancing its pharmacophoric features. The rigid biphenyl and tetrazole ring systems align closely with Losartan’s structure, contributing to high shape and feature similarity.
2. **Valsartan (CID 60846)**
   - **3-D Structural Features**: Valsartan has a similar biphenyl structure with a tetrazole ring. Its additional aliphatic side chain and valine-derived moiety provide slightly different conformations but maintain overall structural similarity to Losartan.
3. **Irbesartan (CID 3749)**

    - **3-D Structural Features**: Irbesartan contains a biphenyl tetrazole core, like Losartan, but features an imidazole ring which adds unique spatial orientation. This contributes to a high degree of 3-D similarity while providing distinct pharmacological properties.
4. **Olmesartan (CID 667490)**
    - **3-D Structural Features**: Olmesartan features a similar biphenyl tetrazole structure but with an additional benzimidazole ring. This complex ring system closely mimics Losartan's 3-D conformation, ensuring significant overlap in pharmacophore features.
5. **Eprosartan (CID 5281035)**
    - **3-D Structural Features**: Eprosartan has a thiophene ring attached to a biphenyl tetrazole structure, differing slightly from Losartan but maintaining key pharmacophoric alignments. The unique orientation of its side chains still falls within the similarity threshold.

These compounds were identified based on their ability to match Losartan's 3-D structural features, such as ring systems and functional groups, which are crucial for their biological activity. The tier 1 approach ensures that these compounds have well-documented biological and pharmacological annotations, enhancing their relevance for further studies.

For more detailed information on these compounds and their structural features, you can explore the PubChem database directly (BioMed Central) (BioMed Central) (Oxford Academic) (BioMed Central)."

### Protocol 4: Getting the Bioactivity Data for the Hit Compounds from Substructure Search

**Snippet of Gold Answer**

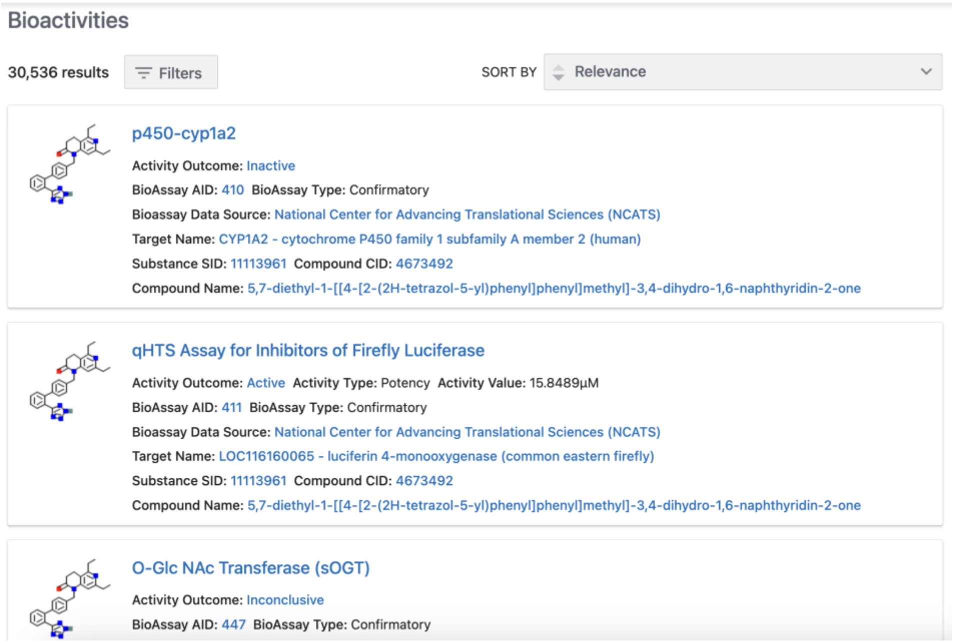
Bioactivities

30,536 results Filters SORT BY Relevance

p450-cyp1a2
Activity Outcome: Inactive
BioAssay AID: 410 BioAssay Type: Confirmatory
Bioassay Data Source: National Center for Advancing Translational Sciences (NCATS)
Target Name: CYP1A2 - cytochrome P450 family 1 subfamily A member 2 (human)
Substance SID: 11113961 Compound CID: 4673492
Compound Name: 5,7-diethyl-1-[[4-[2-(2H-tetrazol-5-yl)phenyl]phenyl]methyl]-3,4-dihydro-1,6-naphthyridin-2-one

qHTS Assay for Inhibitors of Firefly Luciferase
Activity Outcome: Active Activity Type: Potency Activity Value: 15.8489µM
BioAssay AID: 411 BioAssay Type: Confirmatory
Bioassay Data Source: National Center for Advancing Translational Sciences (NCATS)
Target Name: LOC116160065 - luciferin 4-monooxygenase (common eastern firefly)
Substance SID: 11113961 Compound CID: 4673492
Compound Name: 5,7-diethyl-1-[[4-[2-(2H-tetrazol-5-yl)phenyl]phenyl]methyl]-3,4-dihydro-1,6-naphthyridin-2-one

O-Glc NAc Transferase (sOGT)
Activity Outcome: Inconclusive
BioAssay AID: 447 BioAssay Type: Confirmatory

**GPT Generation**

“The bioactivity data for compounds related to the substructure search of 5-(2-phenylphenyl)-2H-tetrazole with the SMILES string C1=CC=C(C=C1)C2=CC=CC=C2C3=N[N]N=N3 can be found on PubChem. Here's a summary of the key information:

1. **AID (Assay Identifier)**: The bioactivity data includes information on various assays used to test the compounds.
2. **Activity Outcomes**: The results of these assays, such as active, inactive, or inconclusive.
3. **Activity Concentrations**: The concentration at which the compound exhibits activity, typically in micromolar (µM) units.
4. **Activity Names**: Names of specific bioactivity assays or targets, e.g., enzyme inhibition, receptor binding.

For more detailed information, including specific assay data and related bioactivity outcomes, you can visit the following links on PubChem and ChemSpider:

- PubChem Bioactivity Data
- ChemSpider Substructure Search"

### Protocol 5. Finding Drugs that Target a Particular Gene

#### Snippet of Gold Answer

| id | cid | dsn | cmpdname | srccmpd | srccmpdname |
|---|---|---|---|---|---|
| 205122 | 2540 | DrugBank | Candesartan Cilexeti | DB00796 | Candesartan cilexeti |
| 362352 | 2541 | DrugBank | Candesartan | DB13919 | Candesartan |
| 417821 | 3749 | DrugBank | Irbesartan | DB01029 | Irbesartan |
| 446240 | 3961 | DrugBank | Losartan | DB00678 | Losartan |
| 1476922 | 60846 | DrugBank | Valsartan | DB00177 | Valsartan |
| 1278296 | 60919 | DrugBank | Tasosartan | DB01349 | Tasosartan |
| 1278299 | 60921 | DrugBank | Saprisartan | DB01347 | Saprisartan |
| 1524664 | 65999 | DrugBank | Telmisartan | DB00966 | Telmisartan |
| 1461907 | 132706 | DrugBank | Forasartan | DB01342 | Forasartan |
| 1382798 | 158781 | DrugBank | Olmesartan | DB00275 | Olmesartan |
| 1412515 | 172198 | DrugBank | angiotensin II | DB11842 | Angiotensin II |
| 1616669 | 5281037 | DrugBank | Eprosartan | DB00876 | Eprosartan |
| 1820297 | 9870652 | DrugBank | Fimasartan | DB09279 | Fimasartan |
| 2144739 | 135409642 | DrugBank | Azilsartan medoxomi | DB08822 | Azilsartan medoxom |

#### GPT Generation

"Based on information from PubChem and related databases, several important drugs interact with the gene encoding the human type-1 angiotensin II receptor (AT1R), the target of losartan. These drugs, known as angiotensin receptor blockers (ARBs), are commonly used to manage conditions such as hypertension, heart failure, and diabetic nephropathy. Here are some of the key ARBs:

1. **Valsartan**: Used to manage hypertension and heart failure. It works by blocking the effects of angiotensin II, reducing blood pressure and improving cardiovascular outcomes (DrugBank) (DrugBank).

2. **Olmesartan**: Effective in treating hypertension, it helps to lower blood pressure and improve heart function by blocking angiotensin II from binding to AT1R (DrugBank).
3. **Telmisartan**: This drug treats hypertension and diabetic nephropathy and is also used to reduce cardiovascular risk. It has a dual role as it also activates peroxisome proliferator-activated receptor gamma (PPARγ), which contributes to its beneficial effects on insulin sensitivity and lipid metabolism (DrugBank).
4. **Irbesartan**: Commonly prescribed for hypertension and diabetic nephropathy. It helps protect the kidneys from damage due to high blood pressure and diabetes (Drugs.com).
5. **Candesartan**: Used to treat hypertension and heart failure, it improves heart function and reduces hospitalizations for heart failure (Drugs.com).
6. **Eprosartan**: This ARB is used to manage hypertension and heart failure, helping to relax blood vessels and lower blood pressure (DrugBank).
7. **Azilsartan medoxomil**: Effective for hypertension, often used alone or in combination with other antihypertensive agents (DrugBank).

These drugs, like losartan, inhibit the action of angiotensin II by blocking its binding to the AT1R, thus helping to control blood pressure and protect against various cardiovascular and renal diseases. For more detailed information on these drugs, you can visit DrugBank or other medical databases (DrugBank) (DrugBank) (Drugs.com)."

### Protocol 6. Getting Bioactivity Data of All Chemicals Tested Against a Protein

**Snippet of Gold Answer**

| baid | activity | aid | sid | cid | geneid | pmid |
|---|---|---|---|---|---|---|
| 99356196 | Active | 39051 | 103213931 | 10373777 | 185 | 2016730 |
| 99371148 | Active | 39666 | 103299812 | 10744859 | 185 | 10893306 |
| 99371149 | Active | 39666 | 103300260 | 10504330 | 185 | 10893306 |
| 99371151 | Active | 39666 | 103300188 | 10393016 | 185 | 10893306 |
| 99371152 | Active | 39666 | 103300252 | 10765881 | 185 | 10893306 |
| 99371153 | Active | 39666 | 103300113 | 11801362 | 185 | 10893306 |
| 99371154 | Active | 39666 | 103300372 | 5249743 | 185 | 10893306 |
| 99371155 | Active | 39666 | 103300688 | 10790819 | 185 | 10893306 |
| 99371157 | Active | 39666 | 103300303 | 10697700 | 185 | 10893306 |

| baid | activity | aid | sid | cid | geneid | pmid | [illegible] |
|---|---|---|---|---|---|---|---|
| 99364399 | Active | 37983 | 103379628 | 44359782 | 81638 | 10579816 | [illegible] |
| 99364401 | Active | 37983 | 160704000 | 172198 | 81638 | 10579816 | [illegible] |
| 99364582 | Active | 37824 | 103379628 | 44359782 | 81638 | 10579816 | [illegible] |
| 99364584 | Active | 37824 | 103379951 | 10558021 | 81638 | 10579816 | [illegible] |
| 99364586 | Active | 37824 | 103546496 | 3961 | 81638 | 10579816 | [illegible] |
| 99364588 | Active | 37824 | 160704000 | 172198 | 81638 | 10579816 | [illegible] |
| 99364599 | Active | 37983 | 103379951 | 10558021 | 81638 | 10579816 | [illegible] |
| 99366829 | Active | 37823 | 160704000 | 172198 | 81638 | 10052967 | [illegible] |
| 99366930 | Active | 37822 | 160704000 | 172198 | 81638 | 10052967 | [illegible] |

### GPT Generation

"Summary of Human Type-1 Angiotensin II Receptor (AT1R)

The human type-1 angiotensin II receptor (AT1R), encoded by the **AGTR1 gene**, is a G protein-coupled receptor (GPCR) that plays a crucial role in the renin-angiotensin system, primarily regulating blood pressure and fluid balance. This receptor mediates various physiological responses, including vasoconstriction, aldosterone synthesis and secretion, increased vasopressin release, cardiac hypertrophy, and modulation of the sympathetic nervous system. AT1R is activated by the peptide hormone angiotensin II (Ang II), which triggers downstream signaling pathways involving G proteins and β-arrestins, leading to diverse cellular responses such as cell proliferation, fibrosis, and inflammation (DrugBank) (Wikipedia).

Bioactivity Data of Key Chemicals Tested Against AT1R

**Human AT1R**

1. **Losartan**: A widely used AT1R antagonist (angiotensin receptor blocker, ARB), effective in treating hypertension and heart failure by preventing angiotensin II from binding to AT1R, thereby reducing vasoconstriction and aldosterone release.
2. **Olmesartan**: Another ARB that binds to AT1R, it is used for managing high blood pressure and has a high affinity for the receptor, providing long-lasting effects.
3. **Candesartan**: Known for its strong binding to AT1R, this ARB helps in reducing blood pressure and improving heart function by blocking angiotensin II effects.

**Rat AT1R**

1. **Valsartan**: Commonly tested in rat models, this ARB blocks AT1R effectively, used to study the effects on blood pressure regulation and cardiac hypertrophy.
2. **Irbesartan**: Similar to other ARBs, it inhibits the action of angiotensin II on AT1R in rats, used to understand its pharmacodynamics and therapeutic potential in cardiovascular diseases.

**Significance in Research**

The AT1R is extensively studied due to its critical role in cardiovascular and renal physiology. Understanding the interactions of various ARBs with AT1R has led to significant advancements in managing hypertension, heart failure, and kidney diseases. Research continues to explore genetic mutations in AT1R and their implications in diseases such as atrial fibrillation, as well as the development of new therapeutic agents targeting this receptor (DrugBank) (Wikipedia) (Frontiers).

For more detailed bioactivity data and further information, please visit PubChem's comprehensive entries on these chemicals and their interactions with AT1R."

### Protocol 7. Finding Compounds Annotated with Classifications or Ontological Terms

#### Snippet of Gold Answer

| cid | cmpdname | cmpdsynonym | mw | mf | polararea | complexity | x |
|---|---|---|---|---|---|---|---|
| 1978 | Acebutolol | acebutolol\|37517-30 | 336.4 | C18H28N2O4 | 87.7 | 401 | |
| 2119 | Alprenolol | alprenolol\|13655-52 | 249.35 | C15H23NO2 | 41.5 | 231 | |
| 2249 | Atenolol | atenolol\|29122-68-7 | 266.34 | C14H22N2O3 | 84.6 | 263 | |
| 2351 | Bepridil | bepridil\|64706-54-3\| | 366.5 | C24H34N2O | 15.7 | 382 | |
| 2475 | Bupranolol | BUPRANOLOL\|Opht | 271.78 | C14H22ClNO2 | 41.5 | 242 | |
| 2583 | Carteolol | CARTEOLOL\|51781 | 292.37 | C16H24N2O3 | 70.6 | 354 | |
| 2663 | Celiprolol | celiprolol\|56980-93- | 379.5 | C20H33N3O4 | 90.9 | 474 | |
| 3333 | Felodipine | felodipine\|72509-76 | 384.2 | C18H19Cl2NO4 | 64.599 | 614 | |
| 3961 | Losartan | losartan\|114798-26- | 422.9 | C22H23ClN6O | 92.5 | 520 | |
| 4171 | Metoprolol | metoprolol\|51384-51 | 267.36 | C15H25NO3 | 50.7 | 215 | |
| 4411 | Solgol | 42200-33-9\|5-[3-(tert | 309.4 | C17H27NO4 | 82 | 344 | |
| 4631 | Oxprenolol | oxprenolol\|6452-71- | 265.35 | C15H23NO3 | 50.7 | 246 | |
| 4946 | Propranolol | propranolol\|Propano | 259.339 | C16H21NO2 | 41.5 | 257 | |
| 5417 | DL-Tetrahydropalma | tetrahydropalmatine\| | 355.4 | C21H25NO4 | 40.2 | 475 | |
| 6100 | Bretylium Tosylate | Bretylium tosylate\|61 | 414.4 | C18H24BrNO3S | 65.6 | 349 | |

Refer to supporting information 7 for the full answer

#### GPT Generation

**"...**

| | **Chemical** | **Therapeutic Uses** |
|---|---|---|
| **0** | **Chemical A** | **['Hypertension', 'Heart Failure']** |
| **1** | **Chemical B** | **['Hypertension', 'Diabetic Nephropathies']** |
| **2** | **Chemical C** | **['Diabetic Nephropathies', 'Left Ventricular Hypertrophy']** |
| **3** | **Chemical D** | **['Heart Failure',** |

| | | 'Hypertension'] |
|---|---|---|

Here is the list of chemicals with therapeutic uses similar to Losartan based on PubChem's MeSH annotations. If you need further details or a different format, please let me know!"

### Protocol 8. Finding Stereoisomers and Isotopomers of a Compound through Identity Search

#### Snippet of Gold Answer

| cid | cmpdname | cmpdsynonym | mw | mf | polararea | complexity | xlogp |
|---|---|---|---|---|---|---|---|
| 60846 | Valsartan | valsartan\|137862-53- | 435.5 | C24H29N5O3 | 112 | 608 | 4.4 |
| 5650 | N-pentanoyl-N-{[2'-(1 | N-pentanoyl-N-{[2'-(1 | 435.5 | C24H29N5O3 | 112 | 608 | 4.4 |
| 5284633 | D-Valsartan | 137862-87-4\|D-Valsa | 435.5 | C24H29N5O3 | 112 | 608 | 4.4 |

Refer to supporting document information 8 part 1 for the full answer

| cid | cmpdname | cmpdsynonym | mw | mf | polararea | complexity | xlogp |
|---|---|---|---|---|---|---|---|
| 60846 | Valsartan | valsartan\|137862-53- | 435.5 | C24H29N5O3 | 112 | 608 | 4.4 |
| 25134333 | (2S)-2-[[dideuterio-[4-[2-(2H-tetrazol-5-yl)ph | | 437.5 | C24H29N5O3 | 112 | 608 | 4.4 |
| 25134334 | (2S)-3-methyl-2-[pentanoyl-[[4-[2,3,4,5-tetr | | 439.5 | C24H29N5O3 | 112 | 608 | 4.4 |
| 25135160 | (2S)-2,3,4,4,4-pentadeuterio-2-[[3-deuterio | | 444.6 | C24H29N5O3 | 112 | 608 | 4.4 |
| 25135222 | Valsartan D9 | Valsartan D9\|Valsarta | 444.6 | C24H29N5O3 | 112 | 608 | 4.4 |
| 25135224 | (2S)-3-methyl-2-[pen | 1089736-75-3\|DTXS | 439.5 | C24H29N5O3 | 112 | 608 | 4.4 |
| 45040680 | Valsartan-d3 | Valsartan-d3\|133190 | 438.5 | C24H29N5O3 | 112 | 608 | 4.4 |
| 87103049 | (2S)-4-deuterio-3-me | SCHEMBL405203 | 436.5 | C24H29N5O3 | 112 | 608 | 4.4 |
| 118856187 | Valsartan-d8 | Valsartan-d8\|108973 | 443.6 | C24H29N5O3 | 112 | 608 | 4.4 |
| 131674069 | (2S)-2,3,4,4,4-pentadeuterio-2-[[4-[2-(2-deu | | 444.6 | C24H29N5O3 | 112 | 608 | 4.4 |
| 156699635 | (2S)-2-[5-deuteriopentanoyl-[[4-[2-(2H-tetra | | 436.5 | C24H29N5O3 | 112 | 608 | 4.4 |

Refer to supporting information 8 part 2 for the full answer

#### GPT Generation

"Based on the information from PubChem and related sources, valsartan (CID 60846) has several stereoisomers and isotopomers. Here are the key details:

##### Stereoisomers of Valsartan

Valsartan has a single stereocenter, and its most commonly known stereoisomer is the (S)-enantiomer. The IUPAC name for this specific stereoisomer is (2S)-3-methyl-2-[pentanoyl-[[4-[2-(2H-tetrazol-5-yl)phenyl]phenyl]methyl]amino]butanoic acid (ChemSpider) (Fisher Scientific).

**Isotopomers of Valsartan**

Isotopomers are isomers that differ only in the isotopic composition of their atoms. For valsartan, common isotopomers include versions labeled with stable isotopes such as deuterium or tritium. Examples include:

- Valsartan-d3
- [$^{3}$H]valsartan (tritiated valsartan) (MacsenLab) (NNLM Guides).

These isotopomers are useful in various research applications, including pharmacokinetics and metabolic studies, because they allow for the tracking and quantification of the drug's behavior in biological systems without altering its chemical properties.

For further details, you can refer to the compound's information on PubChem and ChemSpider."
